# Aromatase-independent estrogenesis: Wood-Ljungdahl pathway likely contributed to the emergence of estrogens in the biosphere

**DOI:** 10.1101/2024.10.24.619986

**Authors:** Po-Hsiang Wang, Tien-Yu Wu, Yi-Lung Chen, Ronnie G. Gicana, Tzong-Huei Lee, Mei-Jou Chen, Tsun-Hsien Hsiao, Mei-Yeh Jade Lu, Yi-Li Lai, Tzi-Yuan Wang, Jeng-Yi Li, Yin-Ru Chiang

## Abstract

Androgen and estrogen, key sex hormones, were long thought to be exclusively produced by vertebrates. The O_2_-dependent aromatase that converts androgen to estrogen (estrogenesis) has never been identified in any prokaryotes. Here, we report the discovery of anaerobic estrogenesis in a Peptococcaceae bacterium (strain TUW77) isolated from the gut of the great blue-spotted mudskipper (*Boleophthalmus pectinirostris*). This strain exhibits unprecedented testosterone fermentation pathways, transforming testosterone into estrogens and androstanediol under anaerobic conditions. Physiological experiments revealed that strain TUW77 grows exclusively on testosterone, utilizing the androgenic C-19 methyl group as both the carbon source and electron donor. The genomic analysis identified three copies of a polycistronic gene cluster, *abeABC* (**<u>a</u>**naerobic **<u>b</u>**acterial **<u>e</u>**strogenesis), encoding components of a classic cobalamin-dependent methyltransferase system. These genes, highly expressed under testosterone-fed conditions, show up to 57% protein identity to the characterized EmtAB from denitrifying *Denitratisoma* spp., known for methylating estrogen into androgen (the reverse reaction). Tiered transcriptomic and proteomic analyses suggest that the removed C-19 methyl group is completely oxidized to CO_2_ *via* the oxidative Wood-Ljungdahl pathway, while the reducing equivalents (NADH) fully reduce remaining testosterone to androstanediol. Consistently, the addition of anthraquinone-2,6-disulfonate, an extracellular electron acceptor, to testosterone-fed TUW77 cultures enabled complete testosterone conversion into estrogen without androstanediol accumulation (anaerobic testosterone oxidation). This discovery of aromatase-independent estrogenesis in anaerobic bacteria suggests that the ancient Wood-Ljungdahl pathway may have contributed to the emergence of estrogens in the early biosphere.

**Significance:** Using a testosterone-grown anaerobic bacterium as a model organism, we characterized this unusual anaerobic estrogenesis at the molecular level. Our findings challenge the long-held belief that estrogen production is exclusive to aromatase-containing vertebrates, expanding our understanding of steroid hormone biosynthesis across domains of life. The involvement of ancient strictly anaerobic Peptococcaceae members and the Wood-Ljungdahl pathway suggests that bacterial estrogenesis may predate O_2_-dependent estrogenesis in vertebrates. Furthermore, the identification of estrogen-producing bacteria in animal guts opens new avenues for potential microbiome-based hypoestrogenism therapies to supplement estrogen in menopausal or ovariectomized females, offering an innovative alternative to current hormone replacement strategies.

## Introduction

Major sex hormones include progestogens, androgens, and estrogens, which modulate the physiology, development, reproduction, and behaviors of animals (1-3). Biosynthesis of C_18_ estrogens from C_19_ androgens proceeds through removing the C-19 angular methyl group, forming an aromatic A-ring (4). This aromatization proceeds through two consecutive hydroxylation reactions of the C-19 methyl group and subsequent oxidative bond cleavage between steroidal C-10 and C-19, which is catalyzed by an aromatase (namely P450arom or CYP19) at the cost of three NADPH and three O_2_ (5) (*SI Appendix*, **Fig.S1**). The reverse reaction (from estrogens to androgens) is thermodynamically challenging; however, the reverse reaction is operated in some denitrifying bacteria. For example, *Denitratisoma* spp. adopt the cobalamin-dependent methyltransferase EmtAB to methylate estrogens to yield 1-dehydrotestosterone (androgenic product with a quinolinic A-ring) (6).

Estrogens were thought to be exclusively produced by aromatase-harboring animals (7, 8). However, our previous data suggested estrogen production through microbial activity in the androgen-contaminated estuarine sediments (9), albeit the microorganisms responsible for anaerobic estrogen production and the underlying mechanisms remained completely unknown. Recently, Jacoby et al. (10) observed the 1-dehydrotestosterone demethylation to form 17β-estradiol (estradiol hereafter) in the *Denitratisoma* cell-free extracts, suggesting that the EmtAB-mediated estrogen methylation is reversible. Although this reverse reaction (transformation of androgen into estrogen) is physiologically irrelevant in the denitrifying *Denitratisoma* spp., the fact that EmtAB could mediate the reverse demethylation of androgens *in vitro* (10) suggests that aromatase-independent estrogen production (estrogenesis) might operate in the anaerobic biosphere. Based on these two lines of evidence, we hypothesized that the observed anaerobic estrogenesis in the estuary sediments could be catalyzed by an EmtAB homolog from unidentified strict anaerobes. This encouraged us to search for microorganisms responsible for the anaerobic estrogenesis. Given that the O_2_ concentration in the estuary sediments is fluctuating, we came up with an idea to enrich the anaerobic androgen-metabolizing microorganisms from the gut of the detritivorous great blue-spotted mudskipper (*Boleophthalmus pectinirostris*) abundantly inhabiting the estuary sediments. The mudskipper gut could provide a stable anaerobic environment for the estrogen-producing microorganisms.

In this study, we isolated and characterized *Phosphitispora* sp. strain TUW77, an extremely fastidious Peptococaceae, from the gut of a great blue-spotted mudskipper. The strain TUW77 exclusively mediates the demethylation of testosterone and uses the androgenic C-19 methyl group as both a carbon source and electron donor. Alternative carbon sources could not be identified under the tested conditions. Physiological experiments indicated that the fermentative strain TUW77 exclusively grows on testosterone, with estrogens and a fully reduced androgen (identified as androstanediol) as extracellular end metabolite. We found that the testosterone demethylation reaction is catalyzed by an EmtAB homolog and a methyl transfer system highly similar to those in methanogens and acetogens, where the androgenic C-19 methyl group is oxidized to yield CO_2_ via the oxidative Wood-Ljungdahl pathway.

## Results

## Results

### A fastidious anaerobe exclusively grows on testosterone

In this study, we first isolated and characterized *Phosphitispora* sp. strain TUW77 (**Fig. 1*A***) from the gut of a great blue-spotted mudskipper (*Boleophthalmus pectinirostris*) (*SI Appendix*, **Fig. S2**). The mudskipper gut microbiota was enriched in an anaerobic chamber using a chemically defined mineral medium (DCB-1) (11) with testosterone (1 mM) as the sole substrate. After 35 days, we observed testosterone consumption (*SI Appendix*, **Fig. S3*A***) accompanied by the production of estradiol (major product), estrone (minor product), and androstanediol (AND). 16S rRNA gene-based analysis revealed enrichment of a Peptococcaceae member, identified as *Phosphitispora* sp. strain TUW77, from ∼0.1% to 46.7% relative abundance. The strain was highly enriched *via* repeated 10^-7^ dilution transfers, yielding a microscopically pure culture of slightly curved, rod-shaped cells in unbranched chains (*SI Appendix*, **Fig. S4**). PCR assay using bacterial universal primers 27F and 1492R (12) (*SI Appendix*, **Table S1**) confirmed a single bacterial 16S rRNA sequence. Phylogenetic analysis showed 98.5% 16S rRNA gene similarity with *Phosphitispora fastidiosa* strain DYL19T (13). Unlike the autolithotrophic strain DYL19T which utilizes inorganic phosphite as the electron donor and CO_2_ as the sole carbon source, organotrophic strain TUW77 grows exclusively on testosterone under fermentative conditions (without exogenous electron acceptor), neither utilizing CO_2_ as the sole carbon source nor oxidizing phosphite (*SI Appendix*, **Table S2**). The bacterial growth experiments also indicated that the strain DYL19T cannot grow on testosterone, highlighting the distinctive metabolic differences between the two *Phosphitispora* spp. Stoichiometric analysis revealed testosterone transformation into three microbial products: estrone, estradiol, and androstanediol under fermentative conditions [**Fig. 1*B***; see *SI Appendix*, **Table S3** for ultraperformance liquid chromatography (UPLC)–high-resolution mass spectrometry (HRMS) patterns of individual products]. After 15 days of anaerobic incubation, ∼2.3 mM testosterone was exhausted, producing 0.9 mM estrogen and 1.4 mM androstandiol (**Fig. 1*B***). Meanwhile, strain TUW77 cell density increased from ∼22,831 cells/mL (16S rRNA gene copies = 91,323 ± 13,785; four 16S rRNA genes/strain TUW77 genome) to 12,927,856 cells/mL (16S rRNA gene copies = 51,711,423 ± 3,278,942) over 15 days, with a doubling time of 52 hours. Most of the microbial products (>90%) were detected in the cell-free supernatant (*SI Appendix*, **Fig. S5**), indicating the production and excretion of estradiol and androstanediol by the testosterone-grown strain TUW77 into the extracellular environment. Furthermore, we did not detect the acetate production in the TUW77 cultures, suggesting that strain TUW77 is not a homoacetogen.

**Figure 1.**
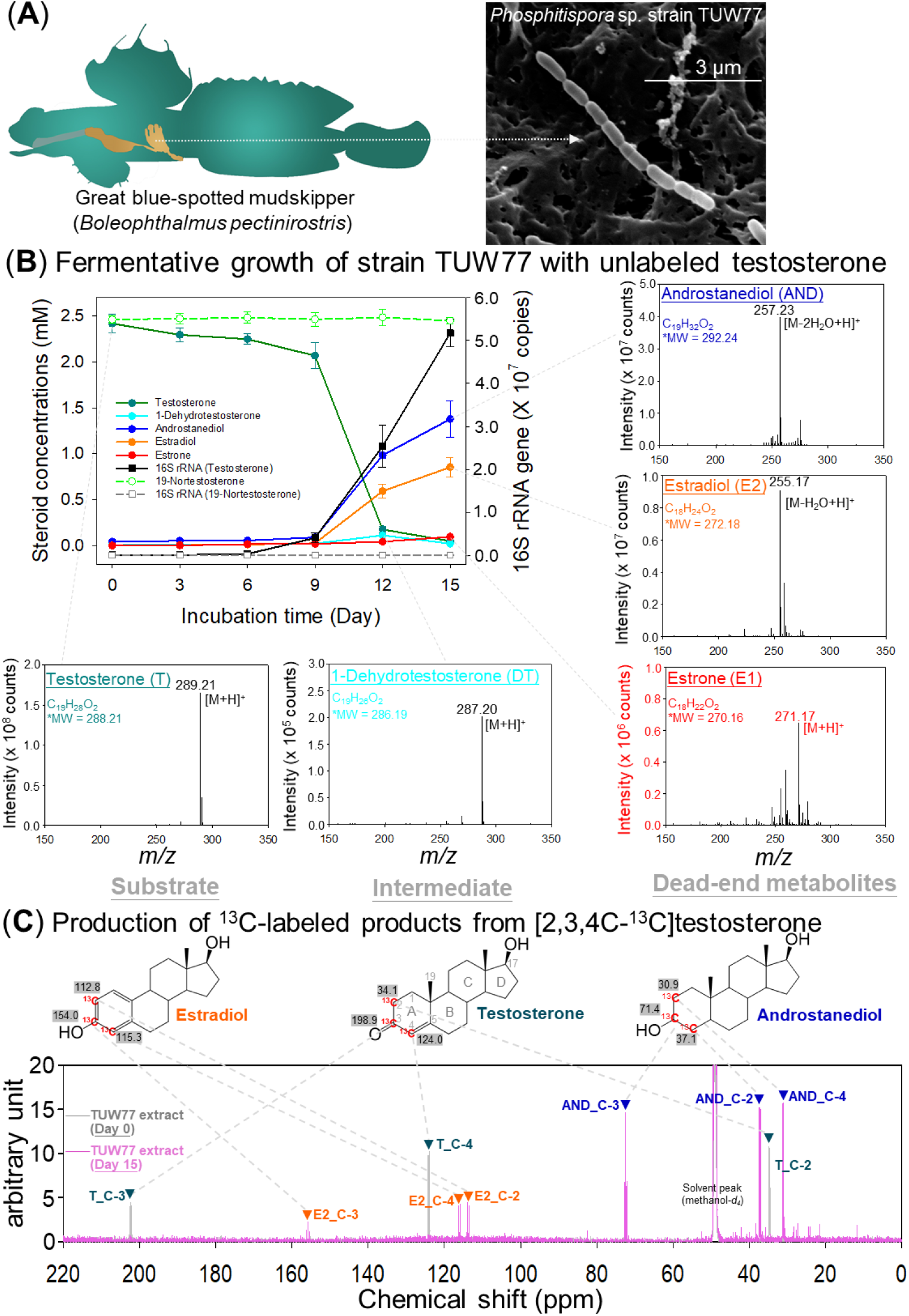
Identification of *Phosphitispora* sp. strain TUW77 and characterization of the anaerobic testosterone metabolism. (**A**) Scanning electron micrograph showing strain TUW77 cells isolated from mudskipper gut. (**B**) Fermentative growth of strain TUW77 with unlabeled testosterone. 19-Nortestosterone served as a reference compound (not metabolized by strain TUW77). Data represent means ± SEM (n = 3). UPLC–APCI– HRMS analysis of the testosterone-grown strain TUW77 culture revealed estradiol and androstanediol as major end products, with 1-dehydrotestosterone as an intermediate. Elemental composition was determined using Mass-Lynx Mass Spectrometry Software (Waters). *, predicted molecular weight (MW) was calculated using the atom mass of ^12^C (12.0000), ^16^O (15.9949), and ^1^H (1.0078). (**C**) ^13^C-NMR spectra of bacterial extracts showing conversion of [2,3,4C-^13^C]testosterone to [2,3,4C-^13^C]estradiol and [2,3,4C-^13^C]androstanediol after 15 days of fermentation. Samples were collected on day 0 (gray) and day 15 (pink), and steroidal metabolites were extracted using ethyl acetate. Chemical shifts (ppm; gray background) for C-2, C-3, and C-4 of ^13^C-labeled steroids were predicted using ChemBioDraw Ultra (version 12.0). Abbreviations: AND, androstanediol; E2, estradiol; T, testosterone.

### An aromatase-independent estrogenesis

Next, we elucidated the mechanisms of anaerobic testosterone fermentation in strain TUW77. The conversion of testosterone into estrogen indicated an androgenic C-10 demethylation reaction that removes the C-19 methyl group from testosterone, yielding C_18_ estrogen (**Fig. 1*C***). This was corroborated by the lack of bacterial growth when the TUW77 cultures were fed with 19-nortestosterone, which lacks the C-19 methyl group (**Fig. 1*B***), demonstrating the crucial role of this methyl group for strain TUW77 growth. To verify the proposed androgenic C-10 demethylation, we analyzed the ethyl acetate extracts of strain TUW77 cultures fed with [2,3,4-^13^C]testosterone using UPLC–HRMS and 13C-NMR spectroscopy. The UPLC–HRMS analysis revealed the dominance of substrate ([2,3,4-^13^C]testosterone) at Day 0 (*SI Appendix*, **Fig. S6*A***). On Day 15, we observed the appearance of three ^13^C-labeled metabolites: [2,3,4-^13^C]androstanediol (**Fig. S6*B***), [2,3,4-^13^C]estradiol (**Fig. S6*C***), and [2,3,4-^13^C]estrone (**Fig. S6*D***). ^13^C-NMR spectroscopy also revealed specific ^13^C-labeling at the C-2, C-3, and C-4 in the major steroidal metabolites (estradiol and androstanediol) (**Fig. 1*C***). The ^13^C-NMR spectra showed doublet peaks for the C-2 and C-4, and triplet peaks for the C-3 in these steroidal metabolites (*SI Appendix*, **Fig. S7**). These UPLC–HRMS and ^13^C-NMR analyses collectively demonstrate testosterone conversion into estrogenic products *via* an unprecedented androgenic C-10 demethylation, likely mediated by an EmtAB homolog (6).

### Tiered OMICs analyses revealed the genes involved in the anaerobic testosterone fermentation

We sequenced and annotated the circular chromosome of strain TUW77 using PacBio HiFi sequencing, yielding a 2,878,163 bp genome with 55.3% G+C content (accession CP156677) (*SI Appendix*, **Dataset S1**). The genome contains four 16S rRNA gene copies (*SI Appendix*, **Appendix S1**) and multiple genes potentially involved in anaerobic bacterial estrogenesis (**Fig. 2*A***). These include four MT1 genes (*abeA*1‒4), four CoP genes (*abeB*1‒4), and three MT2 genes (*abeC*1‒3), with high sequence identity within each gene group. Surprisingly, the deduced amino acid sequences of *abeA* and *abeB* genes show highest similarity (40‒57% identity; *SI Appendix*, **Fig. S8**) to the EmtA and EmtB subunits of estradiol methyltransferase from estrogen-catabolizing *Denitratisoma* spp. (6, 10) (**Fig. 2A**; right panel). The genome also contains a complete set of genes for the oxidative Wood-Ljungdahl pathway, including those encoding the carbon monoxide dehydrogenase/acetyl-CoA synthase complex, CoFeSP complex, and various enzymes in the tetrahydrofolate-carried methyl branch (**Fig. 2B**; *SI Appendix*, **Dataset S1**). Furthermore, the strain TUW77 genome lacks the genes encoding acetate kinase and the AMP-phosphorylating phosphite dehydrogenase (14).

**Figure 2.**
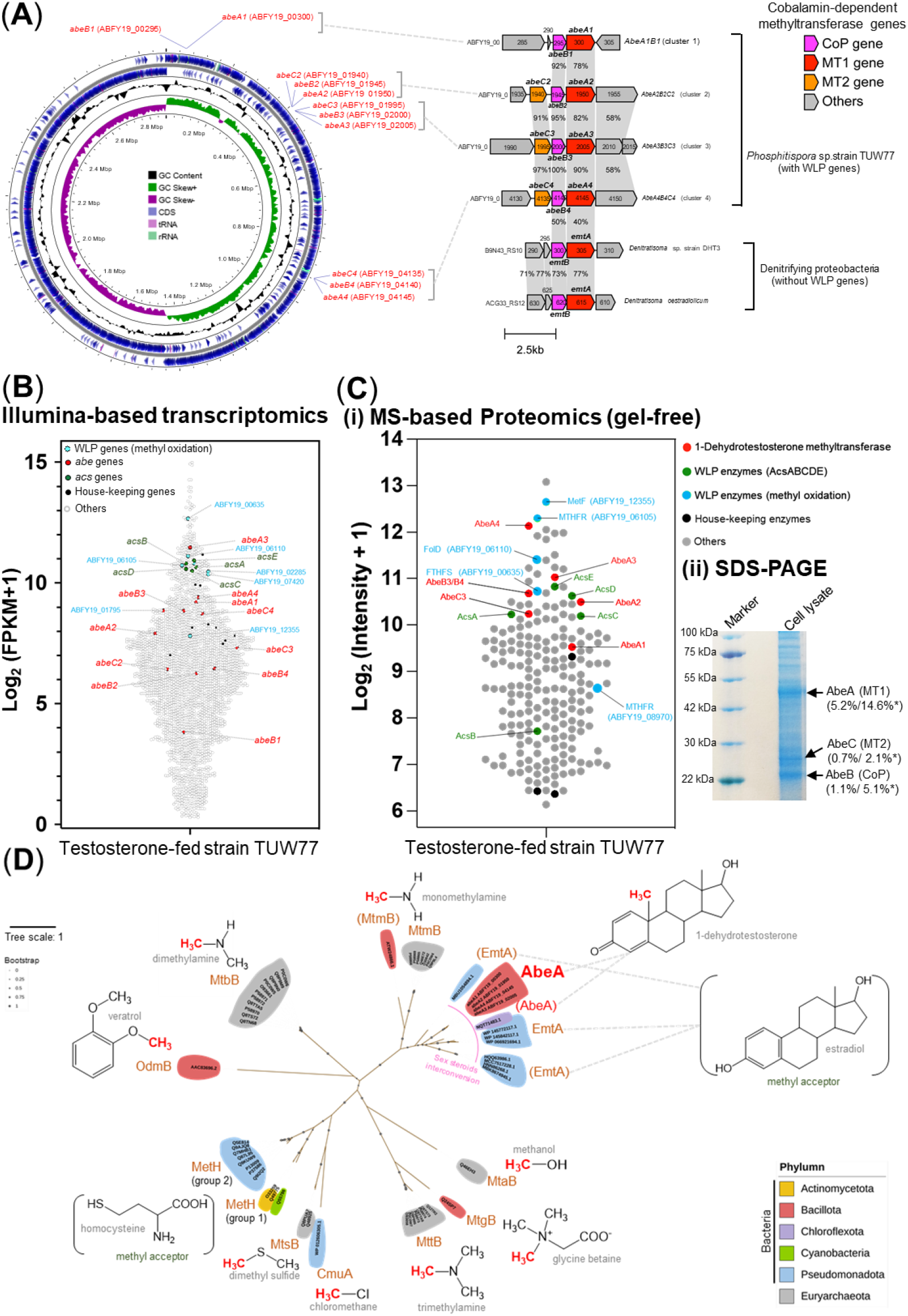
Integrated multi-OMICs analyses reveal the involvement of *abe* genes and corresponding proteins in anaerobic bacterial estrogenesis. (**A**) Comparative genomic analysis of *Phosphitispora* sp. strain TUW77. The paralogous *abe* genes are mapped on the circular chromosome of strain TUW77, with homologous genes identified in testosterone-catabolizing denitrifying proteobacteria *Denitratisoma* spp. Homologous open reading frames (colored arrows) between bacterial genomes are connected by gray blocks. (**B**) Global gene expression profile (RNA-Seq) of strain TUW77 grown anaerobically on testosterone under fermentation conditions. Each spot represents a gene. The FPKM values for genes involved in anaerobic estrogenesis and selected house-keeping genes are provided in **Table S4** and **Dataset S2**. (**C**) MS-based proteomics analysis (**Ci**) and SDS–PAGE (4% to 20%) (**Cii**) of testosterone-grown strain TUW77. *, each Abe protein band contains multiple paralogs; AbeA (5.2% total abundance) comprises AbeA4 (2.7%), AbeA3 (1.2%), AbeA2 (0.9%), and AbeA1 (0.2%). Relative abundances of individual Abe proteins in the soluble protein fraction and corresponding protein bands are listed in **Table S6**. Original gel images are shown in **Fig. S9**; data shown are representative of three independent experiments. (**D**) Phylogenetic analysis of AbeA proteins and characterized cobalamin-dependent methyltransferases (details in **Table S7)**. An unrooted maximum likelihood tree was constructed (bootstrap = 1,000). Parentheses indicate uncharacterized methyltransferases from MAGs.

Strain TUW77 grows exclusively on testosterone, limiting comparative transcriptomic and proteomic investigations with different substrates. We therefore compared expression levels between housekeeping genes (black spots), *abe* genes (red spots), and WLP genes (blue spots) (**Fig. 2*B***). Transcriptomic data reveal significantly higher expression of *abe* genes and WLP genes compared to housekeeping genes during fermentative growth with testosterone (**Fig. 2*B***) [see *SI Appendix*, **Table S4** and **Dataset S2** for gene expression (FPKM+1) values]. Similarly, proteomic data show Abe and WLP enzymes as the most dominant proteins in strain TUW77 proteomes, with many ranking among the top 40 abundant proteins (**Fig. 2*Ci***; *SI Appendix*, **Table S5**). Consistent with these findings, sodium dodecyl sulfate-polyacrylamide gel electrophoresis (SDS–PAGE) of strain TUW77 proteins revealed dominant bands corresponding to AbeA (∼50 kDa), AbeB (∼23 kDa), and AbeC (∼28 kDa), with relative abundances of 5.2%, 1.1%, and 0.7%, respectively (**Fig. 2*Cii***). Subsequent LC–MS/MS analysis of these bands confirmed AbeABC as the most abundant proteins in each band (**Fig. 2*Cii***; *SI Appendix*, **Fig. S9** and **Table S6**). Together, our transcriptomic and proteomic data demonstrate high expression of most Abe and WLP genes and substantial production of AbeABC proteins in strain TUW77 during fermentative growth with testosterone.

### Phylogenetic analysis of *AbeA*

Our data indicated the crucial role of the AbeABC in the aromatase-independent estrogenesis. The MT1 subunit (*e*.*g*., AbeA) of the cobalamin-dependent methyltransferases contains the substrate-binding domain, which mediates the transfer of the methyl group from a specific methyl donor [*e*.*g*., monomethylamine (15) and methanol (16)] to the CoP subunit. Therefore, the MT1 subunit should exhibit the highest substrate specificity among the three subunits. We thus elucidated the phylogenetic relationship of AbeA and other MT1 subunits of the characterized cobalamin-dependent methyltransferases (**Fig. 2D**; *SI Appendix*, **Table S7**). Based on sequence homology, the most AbeA-similar proteins in other microorganisms is the NQT71483.1 protein (identity of protein sequence ∼66%) from the uncultivated Chloroflexota bacterium_bin164 (a marine sediment MAG), while the most similar characterized proteins are the estrogen methyltransferase EmtA (sequence identity ∼40%) from *Denitratisoma* spp. (**Fig. 2*D***).

Moreover, the phylogenetic tree showed that AbeA and EmtA orthologs form a distinct lineage (**Fig. 2*D***), separated from other characterized cobalamin-dependent methyltransferases in prokaryotes. Most unculturable bacteria harboring the AbeA or EmtA are denitrifying proteobacteria (Phylum Pseudomonadota). As the MT2 and WLP are not operated in their genomes, these denitrifying proteobacteria are very likely involved in the retroconversion of estrogen into androgen and the subsequent androgen catabolism. The AbeA/EmtA orthologs were closely placed into the same clade with the monomethylamine methyltransferase MtmB in Firmicutes and methanogenic archaea (*SI Appendix*, **Table S7**), while other bacterial cobalamin-dependent methyltransferases were phylogenetically distant from the AbeA/EmtA orthologs (**Fig. 2*D***).

### Anaerobic testosterone fermentation versus anaerobic testosterone oxidation with anthraquinone-2,6-disulfonate

Our physiological experiments and tiered OMICs analyses indicated that strain TUW77 converts testosterone into estrogen through an androgenic C-10 demethylation, facilitated by the cobalamin-dependent methyltransferase system (AbeABC). Drawing from the established catalytic mechanisms of cobalamin-dependent methyltransferase (17-21) and estrogen methyltransferase EmtAB (6, 10), the process begins with the activation of the testosterone A-ring by a 3-oxo-steroid-1-dehydrogenase (ABFY19_00270), forming a quinonic A-ring (**Fig. 3*A***). Next, AbeA (MT1 component) transfers the androgenic C-19 methyl group from the quinonic A-ring of dehydrotestosterone to the Cob(I) prosthetic group in AbeB (CoP component), resulting in estradiol (C-10 demethylation) (**Fig. 3*A***). Lastly, AbeC (MT2 component) transfers the methyl group from the Cob(III) prosthetic group in AbeB to tetrahydrofolate (THF), entering the oxidative WLP (**Fig. 3*C***). Due to the absence of (***i***) exogenous electron acceptors and (***ii***) acetate kinase for acetogenesis *via* the divergent WLP, the reducing equivalents (NAD(P)H) produced by the oxidative WLP are utilized to fully reduce remaining testosterone to androstanediol (anaerobic testosterone fermentation). To verify this proposed mechanism, we hypothesized that introducing a usable electron acceptor to the testosterone-fed TUW77 cultures would result in complete testosterone conversion into estrogen without androstanediol accumulation. We reviewed the literature for a suitable electron acceptor that could be used by microorganisms gaining energy *via* the oxidative WLP. According to literature, *Methanosarcina acetivorans* can use the humic analog anthraquinone-2,6-disulfonate (AQDS) as an extracellular electron acceptor to oxidize methanol to CO_2_ *via* the oxidative WLP, with a transmembrane multiheme cytochrome C MmcA acting as the conduit for extracellular electron transfer (22). Given the presence of a highly expressed gene (ABFY19_09480) encoding the transmembrane cytochrome C3 in the TUW77 transcriptome (*SI Appendix*, **Figure S10** and **Table S4**), we added AQDS to the testosterone-fed TUW77 cultures as an artificial extracellular electron acceptor. Consistently, this addition resulted in complete testosterone conversion into estrogen without androstanediol accumulation (anaerobic testosterone oxidation) (**Fig. 3*B***). In conclusion, multiple lines of evidence in this study suggest that the C-19 methyl group of testosterone serves as both the electron donor and carbon source for strain TUW77, with estradiol as the end product in bacterial cultures. The reducing equivalents generated during the oxidation of the androgenic C-19 methyl group (*via* the oxidative WLP) are used to reduce the remaining testosterone into androstanediol in a redox-balancing ratio (**Fig. 3*C***).

**Figure 3.**
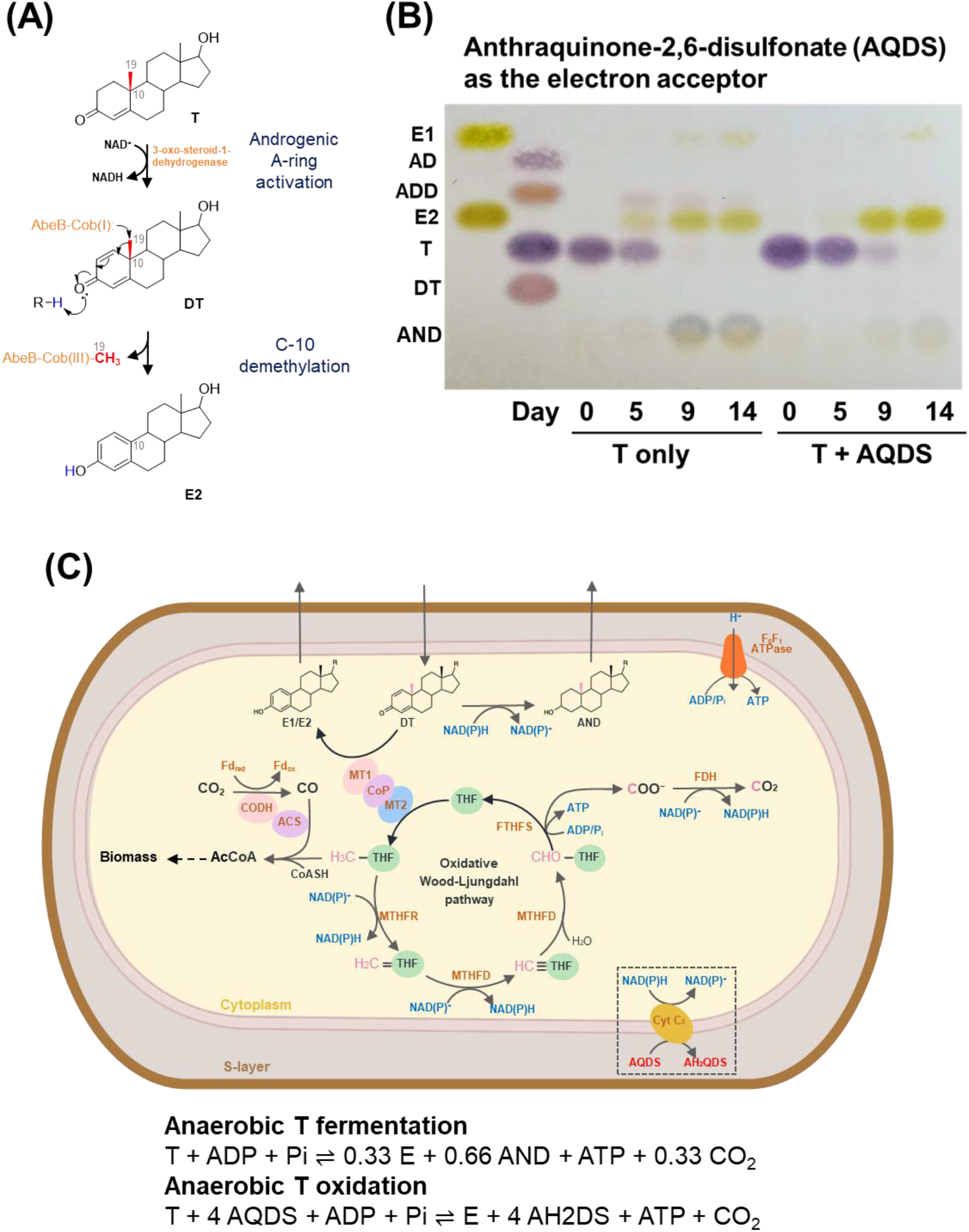
Microbial enzymes, biochemical mechanisms, and stoichiometry of anaerobic estrogenesis. (**A**) Proposed mechanisms operated in the androgenic A-ring activation and Abe-catalyzed, cobalamin-mediated C-10 demethylation. (**B**) TLC analysis showing exclusive estradiol production during anaerobic respiration. Strain TUW77 cells were incubated with and without AQDS for 14 days. The transformation of testosterone into AND occurred only under fermentative conditions. (**C**) Model of metabolic pathways and stoichiometric equations for testosterone fermentation and AQDS-dependent anaerobic respiration. Abbreviations: AD, 4-androstene-3,17-dione; ADD, androsta-1,4-diene-3,17-dione; AND, androstanediol; DT, 1-dehydrotestosterone; E, estrogens (E1 + E2), E1, estrone, estradiol; T, testosterone.

## Discussion

### The anaerobic estrogenesis predated the aromatase-dependent estrogenesis?

In this study, we identified an unprecedented metabolic link between sex steroid biosynthesis and ancient C1 metabolism *via* the AbeABC-mediated androgenic C-10 demethylation. This discovery challenges our understanding of estrogen biosynthesis and its evolutionary history. Sterols (*e*.*g*., cholesterol and phytosterols) are essential membrane components in eukaryotic cells (23). Marine yeasts and microalgae have been proposed as the first sterol producers on earth (24-27). The side-chain removal of sterols through β-oxidation, a common microbial activity, yields C-19 androgens (28-30). Consequently, it is plausible that androgens appeared soon after the emergence of sterols (31, 32).

Prior to our study, the biosynthesis of C_18_ estrogens from C_19_ androgens was thought to be catalyzed exclusively by the O_2_-dependent aromatase present only in vertebrates (8, 33-36). This assumption created a paradox in estrogen receptor evolution: genome mapping and phylogenetic analyses indicate that the most ancient hormone receptors of vertebrates evolved from invertebrate (*e*.*g*., mollusk) steroid receptors (33, 37). According to receptor evolution theory, a new receptor function can only evolve if its ligand is already present. This implies that estrogen should have existed in ancient sediments inhabited by invertebrates, allowing primordial estrogen receptors to access their ligand.

Therefore, our discovery of aromatase-independent estrogenesis in anaerobic bacteria provides a plausible resolution to this paradox. In O_2_- and nitrate-deficient environments, such as estuarine and marine sediments, estrogen is the dead-end product of microbial steroid metabolism. We propose that estrogen produced and excreted by strain TUW77-like bacteria accumulated in ancient sediments over extended periods, facilitating the co-evolution of primordial estrogen receptors and their ligand. Our long-term experiments with mudskipper gut microbiota enrichment cultures also showed that testosterone-derived estrogen remains unaltered over five years (*SI Appendix*, **Fig. S11**). This new understanding of anaerobic estrogenesis not only resolves the evolutionary paradox but also provides fresh insights into the intricate relationship between microbial metabolism and the evolution of complex signaling systems in higher organisms.

### A respiration-like testosterone fermentation process

The testosterone metabolic system observed in strain TUW77 represents an unusual discovery that challenges the contemporary understanding of microbial metabolism, potentially necessitating the reconsideration of current classifications of metabolic processes in biochemical textbooks. This unique process, which we term “respiration-like testosterone fermentation,” defies classification based on current biochemical textbooks, blurring the traditional boundaries between fermentation and respiration. In TUW77, testosterone serves a dual role as both electron donor and acceptor, a characteristic typically associated with fermentation. The C-19 methyl group of testosterone acts as the electron donor, while the steroidal backbone (sterane) functions as the electron acceptor. However, the metabolic pathway exhibits features reminiscent of respiratory processes, particularly in the oxidation of the methyl group via the oxidative WLP and the subsequent reduction of testosterone to androstanediol.

Critically, this process occurs in the absence of exogenous electron acceptors, yet it doesn’t conform to classical fermentation models. The oxidative nature of the WLP in this system, without acetate production, stands in contrast to the divergent WLP observed in heterotrophic homoacetogenesis with CO_2_ as the terminal electron acceptor (38-40). The testosterone metabolism *via* oxidative WLP in TUW77 is more similar to that found in sulfate-reducing bacteria, which use sulfate as a terminal electron acceptor (41-43). The ability of TUW77 to utilize testosterone as an efficient electron sink in anaerobic environments is noteworthy, conferring another role for steroids in microbial energy conservation. This intriguing metabolic strategy may represent an evolutionary intermediate, bridging the gap between strict fermentation and true respiration.

### Ecological and biotechnological implications

The discovery of AbeAB-mediated anaerobic estrogenesis in the strain TUW77 has significant ecological and biotechnological implications. Ecologically, this finding reveals a novel pathway for estrogen production in anaerobic environments like animal guts and sediments, potentially influencing steroid hormone cycling and organismal physiology in these habitats. The presence of AbeAB homologs in uncultivated bacteria from marine sediments suggests this biochemical process may be widespread in anoxic ecosystems. Biotechnologically, this discovery opens new avenues for microbiome-based therapies. For instance, engineered probiotics expressing AbeAB could potentially treat hypoestrogenism in menopausal or ovariectomized females, offering an innovative alternative to current hormone replacement strategies. Additionally, the rarity of *abeAB* genes in microbial genomes makes them, along with 16S rRNA genes, valuable biomarkers for monitoring anaerobic estrogenesis in environmental samples like sediments. This could aid in assessing ecosystem health and tracking steroid pollution. In conclusion, the elucidation of this novel bioprocess expands our understanding of steroid biochemistry and opens up diverse applications in environmental monitoring, bioremediation, and medical treatments.

## Materials and Methods

Strain TUW77 was isolated from the gut of a great blue-spotted mudskipper (*Boleophthalmus pectinirostris*) collected from Guandu estuary, Taiwan (25°6′59.56″N, 121°27′46.99″E). Bacterial cultures were maintained at 30°C in the dark under N_2_/CO_2_ (80:20, v/v) in modified DCB-1 mineral medium with 1 mM testosterone as sole carbon source. For physiological studies, strain TUW77 cells were grown with testosterone under fermentation conditions or with AQDS as terminal electron acceptor for anaerobic respiration. For metabolic studies, strain TUW77 was grown with ^13^C-labeled testosterone under fermentation conditions. Microbial steroid metabolites were analyzed using TLC, HPLC, UPLC–APCI–HRMS, and NMR spectroscopy. Proteins were extracted using B-PER reagents, quantified by BCA assay, and analyzed through SDS-PAGE (4-20%). Label-free quantitative proteomics was performed using LC–MS/MS. The complete genome was sequenced using PacBio HiFi technology. For transcriptomics, rRNA-depleted total RNA was reverse-transcribed and sequenced on an Illumina NovaSeq X Plus. Detailed methods are provided in *SI Appendix*, Supplemental Materials and Methods.

## ACKNOWLEDGMENTS

This study was supported by the Ministry of Science and Technology of Taiwan (109-2221-E-001-002, 109-2811-B-001-513, 110-2311-B-001-033-MY3, and 113-2311-B-001-034-MY3) and Academia Sinica Career Development Award (AS-CDA-110-L13). We gratefully appreciate Ms. Yu-Ching Wu at the Small Molecule Metabolomics Core Facility, Institute of Plant and Microbial Biology, Academia Sinica for the UPLC– HRMS analysis and Dr. Mei-Yeh Lu and her team at the High-Throughput Genomics Core Facility of the Biodiversity Research Center, operated with funding by Academia Sinica Core Facility and Innovative Instrument Project (AS-CFII-108-114).

## DATA AVAILABILITY

The genomic sequences reported in this paper have been deposited in the NCBI GenBank database (accession no.: CP156677).

